# Effects of nystatin on discrete unit 150-3000 Hz extracellular spikes in biofilm forming *Hericium erinaceus* liquid cultures recorded using custom microelectrode arrays

**DOI:** 10.1101/2025.07.16.662945

**Authors:** Davin Browner, Andrew Adamatzky

**Affiliations:** Unconventional Computing Laboratory, UWE, Bristol, U.K

**Keywords:** mycelium, discrete unit potentials, microelectrode arrays, spike sorting

## Abstract

Extracellular electrical potentials have been observed in a number of filamentous mycelial species with incommensurable and non-stereotypic features. In Basidiomycetes, detecting these signals reliably is dependent on the properties of the cell wall and plasma membrane and requires implementation of microelectrode array hardware, filtering and spike sorting methods. In this paper, we present recording methods for detection of discrete unit extracellular spikes in biofilm forming liquid cultures of *Hericium erinaceus*. We utilised custom designed microelectrode arrays (MEAs) with passive planar hard gold microelectrodes and individual radius of 100 *µ*m in recordings at a sample rate of 30 kHz. Triplicate recordings of mycelial samples in a double shielded electromagnetic and RF shield box were conducted for wild-type, ionophore assays and fungicidal assays. The recordings were analysed offline using the Kilosort4 sorting algorithm resulting in detection of discrete unit spikes with milliseconds durations. The clustered spike waveforms for the wild-type triplicates were estimated to have a mean trough-to-peak-time of 2.68 *±* 0.087 ms and width at half maximum of 0.8 *±* 0.031 ms across a combined total of 418 spiking units. Ionophore assays using nystatin solution (10,000 units/ml) exhibited significant statistical differences including a reduction of total units to 97. A decrease in the trough-to-peak time of the mean waveform (1.97 *±* 0.32 ms) and an increase in the width at half maximum (2.7 *±* 2.45 ms) were also observed. Nystatin was found reduce the mean extracellular spike amplitude from 173.06 *µ*V in the wild-type to 25.76 *µ*V in the assay. Physiological disruption of the cell wall and plasma membrane was confirmed by environmental scanning microscopy comparison of triplicates at 90 % humidity. In comparison, a fungicidal assay utilising 12% w/v H_2_O_2_ solution resulted in zero spiking units and absence of discrete unit activity across all channels in triplicate recordings.

## 1. Introduction

Fungal electrophysiology has been studied using different intracellular [1, 2] and extracellular [3] methods since the 1960s. In intracellular studies, potentials can be measured via voltage or current respectively [4]. Using impaled glass microelectrodes, researchers recorded intracellular potentials from *Neurospora crassa*, finding that under standard conditions, the membrane potentials averaged 127 mV, with the inside being negative [1]. These potentials were sensitive to external potassium concentrations and could be quickly reduced by respiratory inhibitors such as sodium azide or the polyene antibiotic nystatin [1]. Subsequent studies explored the relationship between respiration and internal potentials in *N. crassa*. It was discovered that the internal potential has two components: one dependent on respiration and another that persists even under respiratory inhibition [5]. Respiratory inhibitors like azide or 2,4-dinitrophenol were shown to reduce internal potentials from near -200 mV to about -30 mV within minutes. This demonstrated the critical role of respiration in maintaining the electrical properties of fungal cells [5]. Glass microelectrodes used in combination with micromanipulators for fungal cell wall penetration successfully captured spontaneous voltage fluctuations in mycelia of *N. crassa* and exhibited resting membrane potentials of -170 to -230 mV and durations of 1-2 minutes [6]. A cycle of depolarisation and repolarisation of the membrane was observed and was accompanied by a discernible refractory period [6]. In a separate investigation involving *Armillaria bulbosa* mycelial cords, signals were elicited when the expanding mycelium came into contact with a piece of beech wood initially positioned 1–2 cm from the colony [2]. The signals obtained from the cords were assessed using a glass microelectrode inserted among the mycelial strands, with a reference electrode placed in the agar medium. *Pleurotus ostreatus* was also examined using these methods, yielding comparable outcomes in recordings of the periphery of the colony. In both species, the frequency of spontaneous firing was 0.5–5 Hz and amplitude of 5–50 mV [2]. The reported resting potential was between -70 and -100 mV which differs from previous glass microelectrode studies [1]. The rate of the oscillations increased to 20 Hz upon current stimulation.

Patch clamp studies have been conducted for a range of mycelial species and allow for a continuous intracellular environment to be maintained, even after attachment and sealing of the pipette, leading to high precision measurements [7, 8]. The ion channel characteristics of *Phycomyces blakesleeanus*, particularly in relation to osmoregulation and anionic transport have been observed using patch clamp methods. The outward rectifying anionic current, referred to as ORAC, is predominantly activated under hypoosmotic conditions. This current demonstrates outward rectification and bears functional similarities to the vertebrate volume-regulated anion current (VRAC) [9]. The channel is permeable to various anions, including chloride, glutamate, and nitrates [9]. ORAC is a voltage-dependent anion channel with a unitary conductance of 11.3 pS. It is activated by depolarization and exhibits fast activation/deactivation kinetics [10]. The channel exhibits a pronounced selectivity for anions in comparison to cations, as well as a relatively weak selectivity among various anions, adhering to the Eisenman series 1 selectivity sequence [10]. ORAC demonstrates insensitivity to anthracene-9-carboxylic acid while being reversibly inhibited by malate, which suggests a relation to anion accumulation and the regulation of membrane potential [10]. It should be noted that the ORAC patch clamp studies made use of cytoplasmic droplets and not the wild-type mycelium. Studies of *N. crassa* utilising patch clamp techniques have revealed the existence of two distinct types of inositol trisphosphate (IP)-activated calcium channels: a low-conductance channel exhibiting a conductance of 13 pS and a high-conductance channel with a conductance of 77 pS. The low-conductance channel is associated with both the plasma membrane and the endoplasmic reticulum, whereas the high-conductance channel is predominantly linked to vacuolar membranes [11]. The low-conductance channel plays a pivotal role in the establishment of the high calcium ion (Ca^2+^) gradient necessary for effective hyphal growth. Inhibitors targeting this channel, such as 2-aminoethoxydiphenyl borate (2-APB), disrupt the Ca^2+^ gradient and ultimately impedes growth. Additionally, stretch-activated calcium channels have been identified within *N. crassa*, characterised by their permeability to Ca^2+^ and their modulation by intracellular Ca^2+^ concentrations [7, 12].Spontaneous inward potassium (K^+^) channels have been documented in the plasma membrane of Neurospora crassa hyphae. These channels are susceptible to inhibition by tetraethylammonium (TEA), which is recognized as a specific K^+^ channel blocker [7]. The precise role of K^+^ channels in the context of hyphal growth remains underexplored; however, they may play a contributory role in the regulation of membrane potential during the process of tip growth [7]. Anion channels in *Aspergillus niger* display pronounced outward rectification and exhibit selective permeability for chloride ions (Cl) over potassium ions (K). The conductance of a single channel is approximately 43 pS [13]. The patch clamp measurements of the conductance was successful as a result of selective laser ablation of the cell wall fraction at the recording sites. Laser ablation was also utilised to enable *N. crassa* patch clamp recordings [14]. Nanosurgical methods have also been utilised to produce protoplast sections of hyphae and the excised patch currents were predominantly anionic (with selective Cl^-^ making up the largest group of anionic channels detected, followed by non-selective Cl^-^ and organic acids and finally glutamate-permeable) [15]. These studies improved methods for study of protoplasts or cell wall free sections of hyphae in patch clamp as a result of less damaging methods resulting in more physiologically accurate recording conditions [16]. the application of patch clamp methods to mycelium is still complicated by technical challenges in establishing a high-resistance seal, termed a “gigaohm seal” (*≥* 1 GΩ), between the plasma membranes of hyphae and patch clamp while also maintaining realistic physiological conditions.

In extracellular studies, vibrating microelectrode recordings were performed for a number of species in combination with lock-in amplifiers [3, 17, 18, 19, 20]. Electrical currents the water mold *Blastocladiella emersonii* were measured during growth and sporulation. The vibrating probe studies revealed that positive current entered the rhizoid and exited from the thallus in growing cells. The current density was approximately 1 *µ*A/cm^2^, and circumstantial evidence suggested that protons carried much of the current. During sporulation, the current pattern reversed, with positive current entering the thallus and exiting from the rhizoidal region. These findings suggested that electrical currents play a role in the spatial localization of fungal growth and development. Transcellular ion currents in *Achlya* hyphae were studied and the intensity of the current was unaffected by the removal of inorganic constituents from the growth medium but was abolished by an increase in external pH or the deletion of amino acids [18]. Methionine alone diminished the current by two-thirds. The study also revealed a longitudinal pH gradient in the extracellular medium, with the region surrounding the tip being more alkaline and the trunk region being acidic[18]. These findings suggested that a flux of protons, dependent on amino acids, carried the current into the tip and created the surrounding alkaline zone. The proton current was proposed to result from the transport of amino acids rather than their metabolism [18].

Recordings of the extracellular electrophysiology of *Achlya bisexualis, Neurospora crassa, Aspergillus nidulans, Schizophyllum commune* and *Coprinus cinereus* were conducted using vibrating probe techniques [19]. It was observed that in all examined fungi, the current enters at the very tip (apical region, meaning the growing end) and leaves from further back along the hypha [19]. The study measured the magnitude of these transhyphal electrical currents using a vibrating probe technique approximately 30 micrometers from the membrane surface. This allowed for detection of minute DC currents. The current density values ranged between 0.05 and 0.60 picoamperes per square centimeter (pA cm^-2^). These measurements indicate that wider and more rapidly extending hyphae tended to show higher current densities than thinner, slower-growing ones. However, despite these observations, there was no straightforward or linear relationship between the rate of hyphal growth and the current density, highlighting the complexity of the underlying biological processes. Notably, even when the electrical current direction was temporarily reversed during branching in certain fungi, the growth rate was not affected, suggesting that while a correlation exists, the electrical current may not directly dictate growth speed but instead might be involved in other regulatory functions [19]. Later work found that *Neurospora crassa* hyphae drive an electric current through themselves, such that positive charge flows into the anterior and out of distal regions of the trunk [20]. However, the study found similar difficulties in correlating Ca^2+^ to currents. Attempts to detect calcium channels by the use of lanthanide ions and nifedipine were unsuccessful [20]. The galvanotropic behavior of hyphae from *A. nidulans, N. crassa*, and *C. cinerea* was found to be contingent upon both pH and Ca^2+^, indicating a likely role for voltage-gated channels [21]. A change in membrane potential can be observed at the tip of the *N. crassa* during thigmotropic responses [22].

These initial studies have a number of interesting characteristics in addition to the vibrating probe methods. For preparation of *S*.*commune* and *C. cinereus* a different preparation method was used where agar discs were cut from the plates and incubated in a shake culture until pellets with diameters of 15-20 mm were formed [19]. For vibrating probe studies a small sample region was separated from the pellet following 2-3 hours of recovery in a new volume of growth media [19]. High resistivity media was utilised to reduce the number of free ions in the solution with the aim of improving the sensitivity of the measurement instruments. However, by lowering the ion concentration, these media might also alter the natural environment in which the fungal cells operate. The study therefore identified an important trade off between localised ionic noise and attempts to measure common conditions under hyphal and biofilm mediated conduction. The experimental details concentrate on spatial resolution and the distribution of current near the hyphal tips instead of analyzing any alternating or frequency-related behaviors of the potential. The main focus of this work was to measure the DC current density at each point of distance from the apex [19]. Simultaneous recording of different locations was not possible, and the sequential measurement means that if conditions change rapidly, the current at one point might be different by the time another point is measured.

Later studies monitored the longer timescale extracellular electrophysiology using 50 *µ*m tipped electrodes. Extracellular potentials were recorded with the tips contacting aerial mycelium of *S. commune* [23]. The relevant time and frequency domain information was observed in combination with transient voltage values for multiple spatially arranged electrodes using a 24 bit ADC and 1 Hz sample rate. The electrodes were able to detect bioelectrical oscillations from a mass of hyphae. Without pharmacological assays it was difficult to isolate the source of activity and activity was low frequency [23]. The high resistivity of aerial mycelium and the impact on signal conduction were important factors in the resulting signals. Longer timescale biofilm forming submerged liquid cultures of mycelium were recorded using a 50 *µ*m tipped micro-electrode and 100 *µ*m diameter microelectrode arrays to investigate the low frequency and longer timescale activity at 1 Hz sample rates [24]. The low frequency component signals were targeted (0.001-0.5 Hz) and the sample rates precluded detection of high frequency component signals [24]. These bioelectrical oscillations are therefore comparable with similar studies in spontaneous infra-slow or low frequency activity as has been observed in neural tissue [25, 26, 27, 28], epithelial tissue [29] and hydra [30] among other living tissue and organisms.

In this paper, we outline results relating to analysis of high frequency component and discrete unit potentials in the species *Hericium erinaceus* recorded and processed using a custom MEA in combination with offline spike sorting algorithm. Sample rates of 30 kHz and microelectrodes with radius of 100 *µ*m achieved recordings with detectable discrete unit extracellular spikes that were preserved upon bandpass filtering of 150-3000 Hz. We identified discrete unit spikes exhibiting stereotypical waveforms with troughs and post-trough peaks. The resulting waveforms were analysed and had a mean trough-to-peak-time of 2.68 *±* 0.08 ms across a total of 418 spiking units for all wild-type triplicates. The nystatin assays showed significant statistical differences in the resulting spike waveforms including a reduction of total units to 97, increase in the trough-to-peak time of the mean waveform (1.97 *±* 0.32 ms) and in the width at half maximum (2.7 *±* 2.45 ms). The magnitude of extracellular spikes reduced from 158.76 *µ*V to 25.76 *µ*V. Physiological disruption, via lipid peroxidation, of the cell wall and plasma membrane was confirmed by environmental scanning microscopy based comparison of triplicates at 90 % humidity. The multicellular structure, thick cell walls, biofilm formation and lower permeability likely make this species highly tolerant or largely resistant to nystatin. However, ion leakage, stress and disrupted signaling can still occur in the absence of fungicidal effects. As a result, nystatin appears to have a specific role as a modulator of the properties of the spike leading to changes in the mean waveform morphology as recorded here via extracellular microelectrodes. A fungicidal assay utilising 12 % w/w solution of H_2_O_2_ which resulted in zero sorted spiking units.

## 2. Methods

### Biofilm forming static submerged liquid cultures

The static submerged liquid cultures of *H. Erinaceus* were grown from stocks of agar plate samples in malt liquid culture media (1.5 %). Aeration was limited to a single syringe Filter (13mm diameter and 0.22 *µ*m filter size). No stirring or aeration was utilised and the static submerged liquid fermentation methods resulted in the formation of biofilms in the liquid culture.

### Micro-electrode arrays (MEAs)

The microelectrode array (MEA) was a custom printed circuit board (PCB) with 10U” hard gold coating. The MEA had 64 channels including a reference electrode. The single ended circular electrodes in the MEA are arranged in a rectangular grid array with each electrode having a radius of 100 *µ*m and a vertical and horizontal spacing of 700 *µ*m. The exposed electrically conductive surface of the electrodes was equivalent to the diameter at 200 *µ*m each. Reference electrodes and channel locations are depicted in Figure 1 (B). Recordings were conducted in a shielding box (Tescom, South Korea). The shielding box functioned as a Faraday cage with copper mesh lining and also blocked high frequency noise. With RF shielding effectiveness of up to *>* 80 dB across a wide frequency range, the TC-5910D also acted as a Faraday cage as a result of its conductive copper mesh lining, steel exterior and RF-filtered connectors resulting in a fully enclosed and dark recording environment. The shielding box was placed on an anti-vibration table (Adam Equipment, U.K.) to eliminate mechanical vibration artifacts. Low frequency components and line noise were removed by the bandpass (150-3000 Hz) and common average reference procedures. The effect of liquid media in the array chamber was minimised by dehydration using a q-tip placed gently above the in-situ culture for 10 seconds. Mycelial samples were rested for 2 hours in-situ to recover from transfer shock.

**Figure 1:**
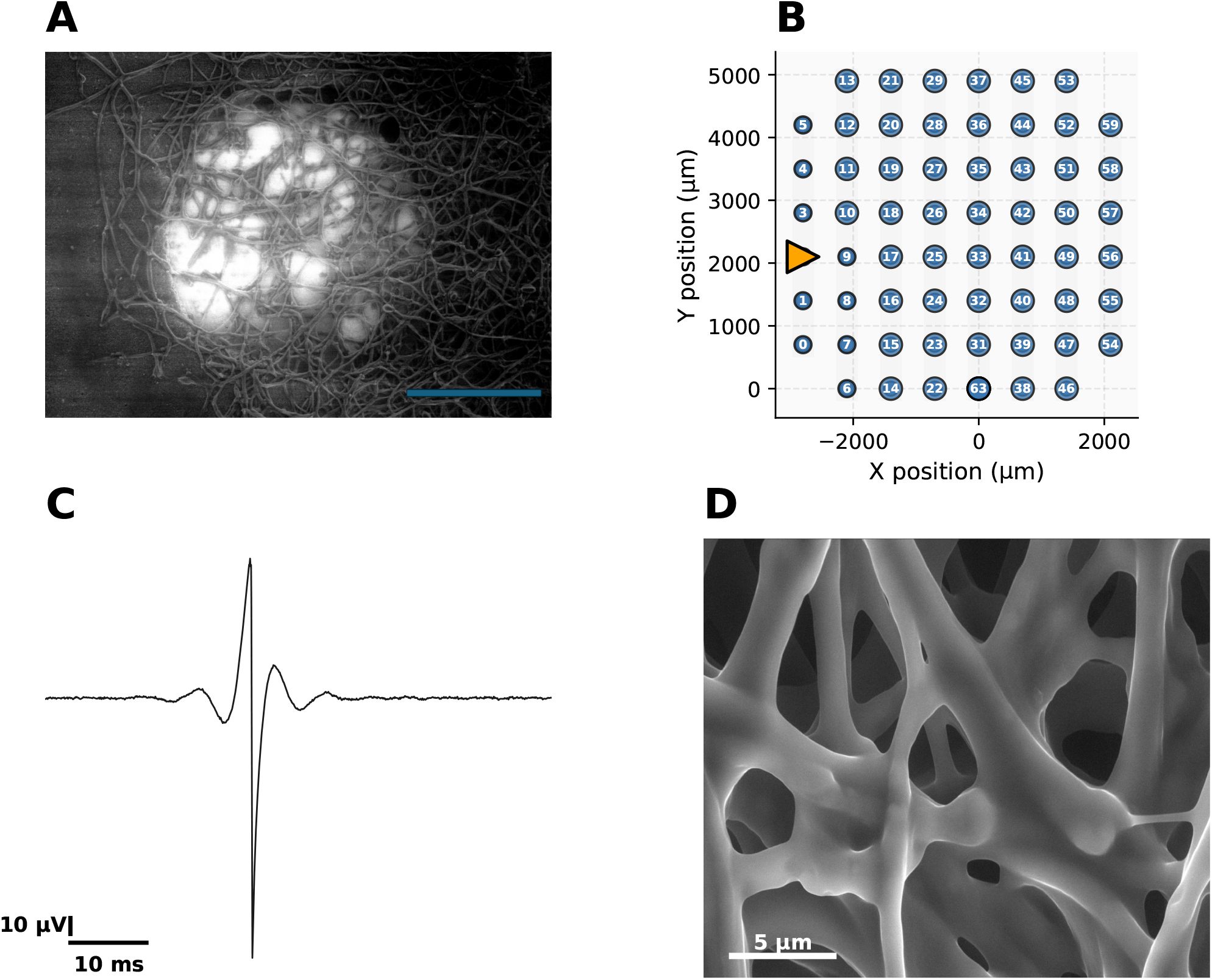
(A) Micrograph of the EPS containing liquid mycelium (100 *µ*m scalebar, magnification of 534, Hv of 10 kV, internal pressure of 4 Torr and HFW of 388 *µ*m);(B) Layout of the MEA used in this study including numbered microelectrodes and reference electrode (orange triangle);(C) Single spike from a channel trace showing the discrete unit mycelial potential;(D) Micrograph of biofilm forming liquid cultures showing kayers of EPS on the surface of hyphae in a hydrated sample (5 *µ*m scalebar, magnification of *×*6906, Hv of 9.50 kV, internal pressure of 5 Torr, HFW of 30 *µ*m and imaged at 90 %) humidity.

### Amplification

The Intan RHD2164 (Intan Technologies, U.S.) was used to amplify the signals from the MEA. It is a 64-channel digital electrophysiology interface chip designed for high-density extracellular recordings. Features 64 low-noise amplifiers with an input-referred noise of 2.4 *µ*V RMS and an input impedance of 1 GΩ // 2 pF. The bandwidth is programmable, with a low cutoff frequency ranging from 0.1 Hz to 500 Hz and a high cutoff frequency from 100 Hz to 20 kHz. The chip includes a 16-bit analog-to-digital converter (ADC) that provides simultaneous sampling of all channels at a maximum sampling rate of 30 kS*/*s per channel. Communication is facilitated through an SPI interface. The MEA was connected via custom housing (connection junctions had a maximum length of 150mm) and routed to the amplifier using an electrode adapter board (Intan Technologies, U.S.). The fungal sample, MEA, and Intan headstages was placed inside the shielding box. The acquisition board controller and computer (which generate digital noise and have high-voltage AC power wiring) were located outside the shielding box. The GND terminal on the Intan headstage was connected to the tissue via a ground reference electrode. The zero-ohm R0 resistor was left in place on the RHD headstage so that REF could be shorted to GND. This resulted in a combined REF/GND electrode that was connected to the fungal sample. The fungal sample was electrically isolated from the shielding box. All recordings were conducted in a temperature controlled environment to minimise thermal noise (19^*°*^C). A butyl rubber lid covered the recording chamber to maintain humidity levels of 90-95 %. Photoelectric noise was negligible due to the dark environment of the interior of the shielding box. The shielding box was locked shut during recordings and was only opened for inspection or changing of samples.

### Data acquisition

The Open Ephys Acquisition Board (OpenEphys, Portugal) was used for data acquisition. It is an open-source interface designed for high-channel-count electrophysiology experiments [31]. In our experiments, a single Intan RHD2164 headstage was connected to the acquisition board via SPI cable. This completed the recording setup and allowed for simultaneously sampling at a per-channel raw recording frequency range of 0.1 to 6500 Hz used in experiments.

### Spike sorting

Recordings had a 0.1-6500 Hz bandpass filter implemented using the analog filters of the aquisition device during recordings. Following acquisition, a digital bandpass filter was used to isolate activity between 150-3000 Hz using a Butterworth second order filter. This frequency range was selected as a result of power spectral density analysis indicating an absence of frequency components *≥* 3000 Hz. A common reference electrode was used to reduce non-physiological noise further by averaging and common mode noise rejection. Further analysis and spike sorting used the spikeinterface library [32]. The Kilosort4 sorting algorithm was used to detect and sort spikes [33]. Kilosort4 is a high-performance spike sorting algorithm designed for electrophysiological data, specifically for analyzing extracellular recordings. It operates using a multi-step pipeline that includes pre-processing, spike detection, clustering, and post-processing. Following bandpass filtering the data was whitened to reduce correlations and improve detection. Candidate spike events were then detected. The spatiotemporal features of the detected spikes were extracted and matched to the MEA layout. Clustering was performed with templates representing the average waveform of each cluster generated. Noise clusters and artifacts were removed based on statistical criteria.

### Ionophore and fungicidal assays

Ionophore assays were prepared to assess if there was a resulting change in the discrete unit waveforms. Nystatin, the polyene antifungal drug, was utilised because it exhibits a broad antifungal spectrum and acts on sensitive organisms by destroying the selective permeability of the plasma membrane by pore formation in surrounding tissue and lipids [34]. It was shown to cause a general leakage of small molecules (amino acids, nucleotides, as well as K^+^) from yeast cells [35, 36, 37]. Here, a 200 *µ*l aliquot of nystatin suspension, 10,000 unit/mL in Dulbecco’s Phosphate-Buffered Saline (DPBS) (Sigma Aldrich, U.S.) was injected into the growing cultures and left to permeate for 2 hours with recordings of 1 hour for each of the triplicates. Following this, all measurements of the triplicates were conducted in identical conditions to the non-assayed cultures. Leakage of cell contents was confirmed in the assayed samples (see Figure 1 (D) and). The resulting traces from the ionophore assay were compared with a fungicidal assay comprising of a 200 *µ*l aliquot of the 12 % H_2_O_2_. The fungicidal assay was left for 2 hours in solution and was confirmed to be non-viable via subsequent transfer to an agar plate. 12 % H_2_O_2_ was found to be an effective fungicidal assay for these samples and even when containing EPS materials.

### Environmental scanning electron microscopy

Environmental scanning electron microscopy (ESEM) allows the imaging of fully hydrated biological samples without traditional SEM preparation involving dehydration and coating with thin electrically conductive outer layers. The FEI Quanta 650 FEG scanning electron microscope (SEM) (FEI Company, U.S.) was used to image the samples in a hydrated state and produce micrographs. The hydrated samples were fixed in 2% glutaraldehyde in PBS for 1 h at room temperature and then rinsed (3 × 1 h) in deionised water and stored at 4 until required. Biofilm forming liquid cultures were placed on a stainless steel stub on the Peltier stage that was set to cool to 2^*°*^C. At this temperature, the dew point of water was lowered below 10 Torr. A drop of distilled water was placed in each of the four depressions in the Peltier substage to promote rapid increase in humidity. The chamber was pumped to 7.5 Torr. The set pressure was then reduced in 0.2 Torr steps to 6.7 Torr. The surface of the water drop was observed, and the water evaporated slowly by reducing the pressure to expose the fully hydrated samples. A repeated humidity modification was used for each sample dropping to 90 % and remaining at this humidity level for the duration of imaging.

## 3. Results

Scanning electron micrographs of the biofilm forming liquid mycelium were imaged using ESEM at a constant humidity of 90 % preserving EPS components. The EPS containing mycelial samples are shown in a micrograph in Figure 1 (A) growing in-situ on the microelectrode array and (D) following electrophysiological recording with the EPS constituents preserved. EPS in the wild-type are shown in micrographs in Figure 2 (A) - (C). In contrast Figure 2 (D) - (F) shows the nystatin assayed cultures with evidence of cellular leakage and disruption of the cell wall. In these samples the EPS also appeared to be disrupted.

**Figure 2:**
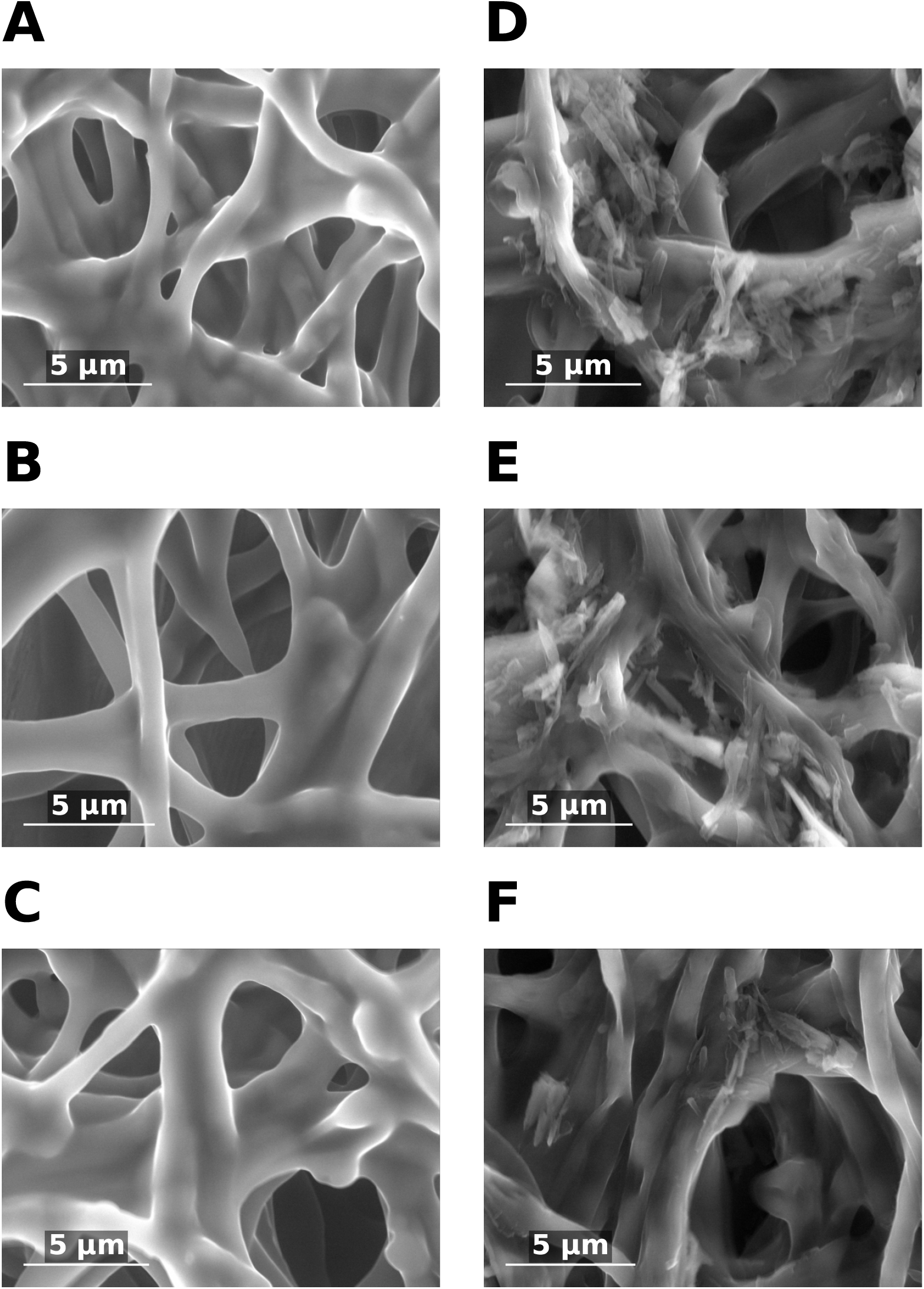
A) Triplicate 1: *H. erinaceus* biofilm (magnification of x6906, Hv of 9.50 kV, internal pressure of 5 Torr and HFW of 30 *µ*m);(B) Triplicate 2: *H. erinaceus* biofilm (magnification of x8956, Hv of 9.50 kV, internal pressure of Torr and HFW of 23.1 *µ*m);(C) Triplicate 3: *H. erinaceus* biofilm (magnification of x8447, Hv of 9.50 kV, internal pressure of 5 Torr and HFW of 24.5 *µ*m);(D) Triplicate 4 - nystatin assay (10,000 units/ml) (magnification of x8336, Hv of 10.00 kV, internal pressure of 4 Torr and HFW of 24.9 *µ*m);(E) Triplicate 5 - nystatin assay (10,000 units/ml)(magnification of x8336, Hv of 10.00 kV, internal pressure of 4 Torr and HFW of 24.9 *µ*m);(F) Triplicate 6 - nystatin assay (10,000 units/ml)(magnification of x8336, Hv of 10.00 kV, internal pressure of 4 Torr and HFW of 24.9 *µ*m).

The detection of discrete unit spikes and these associated features can be facilitated by use of medium sized microelectrodes (e.g. with radius of 100 *µ*m). In microscopy of the wild-type samples, septate junctions were observed c. every 50-100 *µ*ms. In Basidiomycota the exterior of the hyphae consists of a cell wall and a semi-permeable bilayer cell membrane called the plasma membrane. In general, hyphae of the lower fungi, i.e. the Glomeromycota, Zygomycota, and Chytridiomycota are sparsely, if at all, septated resulting in a continuous structure [38]. Hyphae of the higher fungi, i.e. the Ascomycota and Basidiomycota, are compartmentalised by septa [38]. These septa contain central pores of up to 500 *nm* that allow streaming of cytoplasm and translocation of organelles like mitochondria and nuclei [38]. Gap junctions in animals and plasmodesmata in plants have pores with a diameter of about 1.5 to 3.0 *nm*. However, the the diameter of the pores of plasmodesmata and gap junctions is dynamic [38]. Heterogenity of the studied hyphae should be considered in extracellular electrophysiology when selecting the spatial dimensions of sorting algorithms as well as microelectrode sizes and layout. If the plasma membrane exhibits a fully continuous structure enabling cytoplasmic and organelle flow among hyphal compartments. This structure would allow for propagation of electrical signals in an insulated cable-like structure with less internal electrical resistance [39]. However, hyphal networks punctuated by septal pores at intervals along the extent of individual hypha could have different electrophysiological characteristics when recorded from extracellular environment [39].

The plasma membrane and cell wall are also implicated in the transmission of intracellular potentials to the extracellular environment. In general, the plasma membrane and cell wall separates different ion concentrations on the inner and outer sides of the membrane. The preservation of resting membrane potentials is achieved actively within the cell through the regulation of ion movements across the membrane, facilitated by selective ion channels and/or pumps [40]. In active rather than passive ion channels the concentrations are also represented in terms of their charges and the resulting electrochemical gradient forms a membrane potential. In theory and in stereotypical conditions, electrophysiological changes are due to opening ion channels due to chemical or electrical stimulation and the corresponding ions move along a transport dependent and localised electrochemical gradient. The resistance of the membrane is lowered resulting in an inward or outward flow of ions. When measured using intracellular methods this is represented by the transmembrane current. The extracellular fluid is conductive and typically exhibits a low resistance that is not nil. The extracellular current results in a small voltage magnitude that can be measured with proximate electrodes localised to the target tissue. Its current is dependent on Ohm’s law (*U* = *R ∗ I*) and is dependent on the properties of the tissue, its relevant active and passive transport phenomena and the operation of voltage and chemical gated ion channels. The extracellular signals are smaller than the transmembrane potentials and this depends on the distance of the source to the electrode.

Extracellular recordings exhibit signal amplitudes that decrease with increasing distance of the electrode from the signal source. A close interface between the electrode and the cell membrane produces a high signal-to-noise ratio. Typically, distances over 100 *µ*m from an idealised single target cell lead to noise from diffusion phenomena and/or the activity of nearby cells. However, this cannot be applied liberally to all biological systems due to differences in mechanical and electrical properties of tissue as well as the dynamic composition of the extracellular matrix of different cells and organisms. Both the transmembrane current and the extracellular potential follow the same time course, with minor differences due to noise, and are roughly equivalent to the first derivative of the transmembrane potential. If the signal is present in the intracellular environment and is of a high enough magnitude then it should be detectable in the proximal region to the target hyphae provided suitable EPS or other electrically or ionically conductive substance is present. Agar was found to be a poor intermediary of such signals due to the introduction of electrochemical noise and non-physiological electrical potential shifts such as Donnan potentials. As a result, malt liquid culture media (1.5 %) was preferred in cultivation. The layout of the MEA used in triplicate recordings is shown in Figure 1 (B) and comprised of 64 hard gold microelectrodes including one reference electrode that was connected to GND to remove common mode noise. The reference electrode (triangular shaped orange electrode) is highlighted to distinguish it from the single ended recording electrodes.

Discrete unit spikes can be defined as having transients of milliseconds duration with frequency components above 150 Hz. For instance, visible spikes following bandpass filtering (e.g. 150-3000 Hz). In extracellular recordings with suitable proximal microelectrodes discrete unit activity should appear as rapid troughs followed by a return to the baseline peak. A further peak prior to the trough may be apparent. A single isolated spike with a duration of c. 8 ms from the MEA is shown in Figure 1 (C) to illustrate the characteristics of the discrete unit activity. The initial peak, trough and peak after trough can be observed in the isolated spike.

Mycelial discrete unit spikes were confirmed for the EPS containing triplicate wild-type *H. erinaceus* cultures visually in single trace and all channel activity maps prior to sorting. Prior to bandpass filtering spikes in the range of 0.1-6500 Hz were evident in the wild-type samples. These full bandwidth spikes can be seen in the 300 s single channel trace in Figure 3 (A)-(C). In contrast the nystatin assayed cultures were dominated by local field potential-like oscillations of a lower magnitude in the 0.1-6500 Hz range. A 300 s single channel trace of the nystatin assays can be seen in Figure 3 (D)-(F). Following bandpass filtering (150-3000 Hz) and common average referencing (CAR) the discrete unit spikes in the wild-type samples were preserved as shown in single channel trace plots with durations of 300 s in Figure 3 (A)-(C). Local field potential-like oscillations in the nystatin assays were largely absent following bandpass and CAR filtering. A 300 s sample of the triplicate assays can be seen in Figure 3 (D)-(F). Comparison of the traces shows a significant visual differences between (0.1-6500 Hz) and the bandpass filtered (150-3000 Hz) recordings including a reduction of low frequency noise. Traces from each triplicate recording for all channels of the MEA post bandpass and CAR filtering are plotted in map mode in Figure 3 (A)-(F) with negative or positive biased peaks indicated by cmap color gradients as denoted in the colourbar.

**Figure 3:**
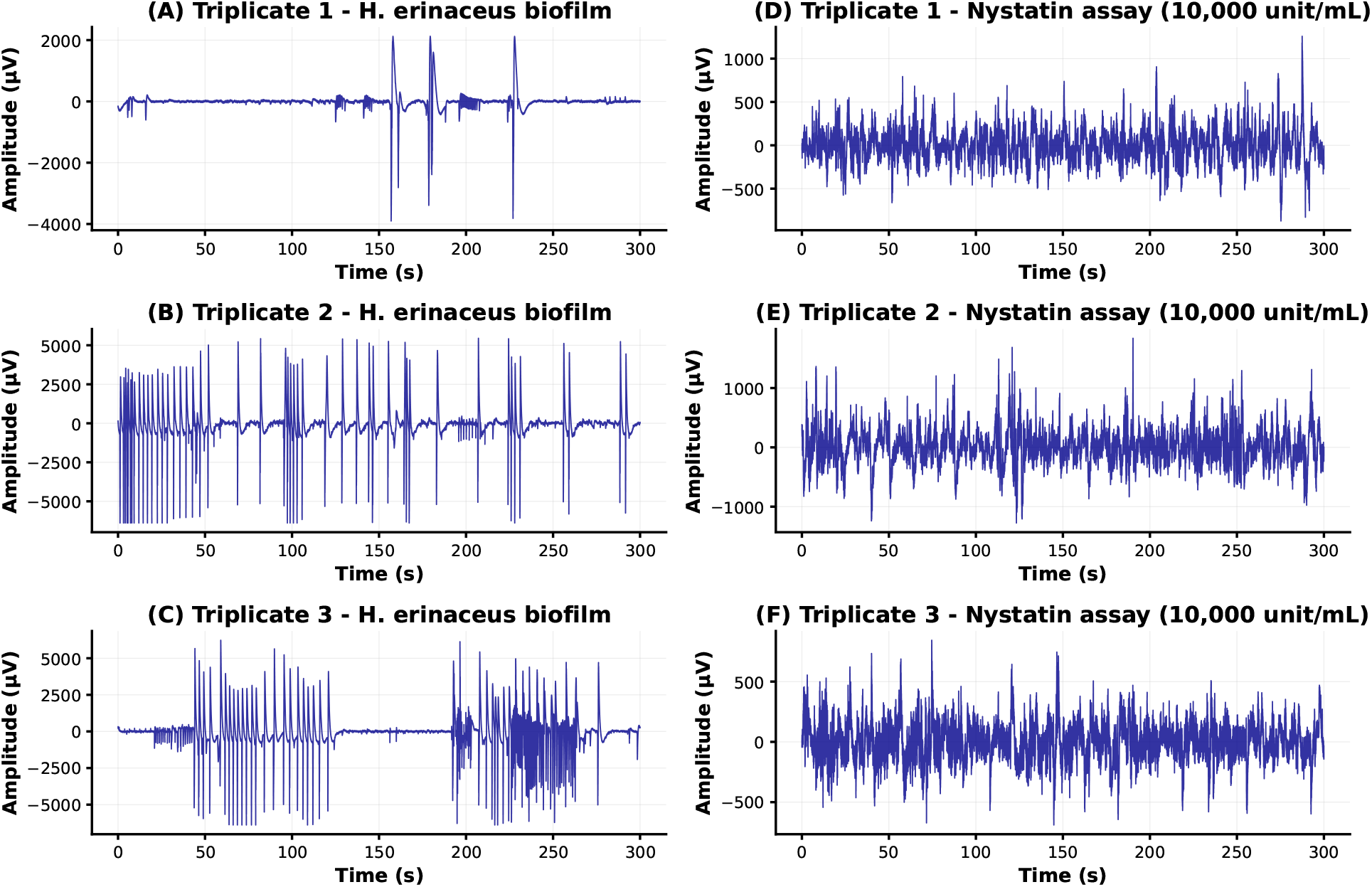
Raw signal traces (0.1-6500 Hz): (A) Triplicate 1: *H. erinaceus* biofilm;(B) Triplicate 2: *H. erinaceus* biofilm;(C) Triplicate 3: *H. erinaceus* biofilm;(D) Triplicate 4 - nystatin assay (10,000 units/ml);(E) Triplicate 5 - nystatin assay (10,000 units/ml);(F) Triplicate 6 - nystatin assay (10,000 units/ml).

**Figure 4:**
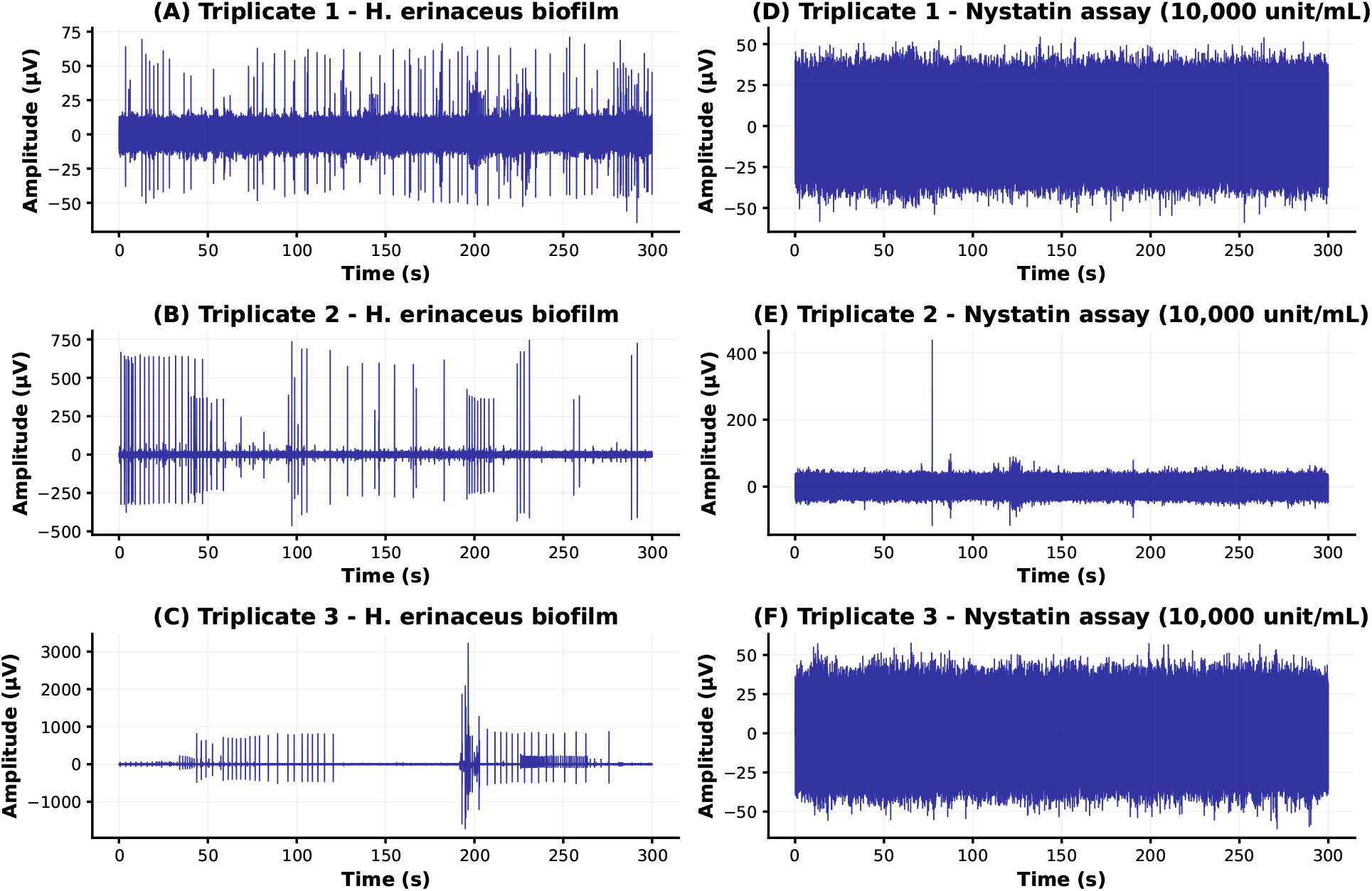
Filtered signal traces (150-3000 Hz): (A) Triplicate 1: *H. erinaceus* biofilm;(B) Triplicate 2: *H. erinaceus* biofilm;(C) Triplicate 3: *H. erinaceus* biofilm;(D) Triplicate 4 - nystatin assay (10,000 units/ml);(E) Triplicate 5 - nystatin assay (10,000 units/ml;(F) Triplicate 6 - nystatin assay (10,000 units/ml).

We performed spike sorting on the filtered recordings using Kilosort4. Modification of the sorting parameters was required to parse the discrete unit mycelial spikes. Thresholds were increased to *Th*_*u*_*niversal* = 12 and *Th*_*l*_*earned* = 13. Drift correction was disabled due to the lower channel count of the MEA (*nblocks* = 0). The number of time samples to represent the spike waveforms was increased to 9 times the default setting (*nt*) to account for the anticipated longer timescale of mycelial spikes. A bandpass filter of 150-3000 Hz was implemented to match the filters of the processed recordings.

The discrete unit waveform analysis revealed distinctive characteristics for both conditions. The mean magnitude of extracellular spikes (i.e. the mean amplitude of the trough) reduced from 173.06 *µ*V *±*353.26 *µ*V in the wild-type compared with 25.76 *µ*V *±*22.59 *µ*V in the nystatin assay indicating that significant disruption of the EPS and cell wall occurred. The mean firing rate for the wild-type cultures was 1.07 *±* 2.32 Hz compared with 8.99 *±* 20.20 Hz in the nystatin assay. Inter-spike intervals were lower in the nystatin assay 106.38 ms compared to the wild-type 459.10 ms. These statistical measures are plotted in Figure 6 for the wild-type and Figure 7 for the nystatin assay. The large standard deviation and low magnitude of spike values in the nystatin assay suggests atypical spiking patterns that may be contaminated by electrochemical noise as a result of pore formation and cell leakage. Nystatin is primarily known as an ionophore that forms pores in the fungal cell membrane via binding to ergosterol. This leads to K^+^ leakage, acidification and the death of the fungus in sufficient concentrations. Filamentous basidiomycetes may be less susceptible due to structural and delivery barriers compared with yeasts.

**Figure 5:**
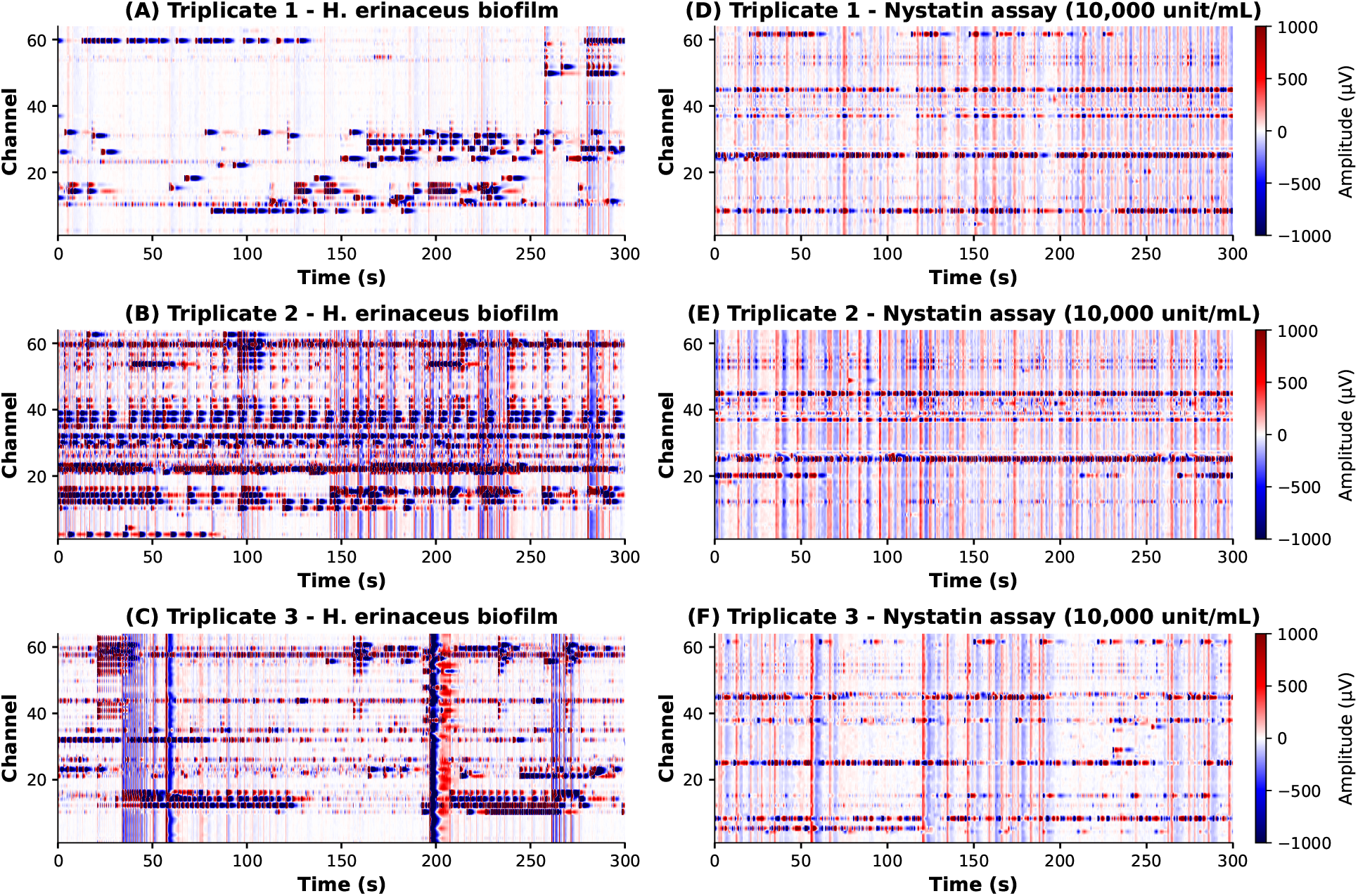
Bandpass and CAR filtered channels trace map (150-3000 Hz) 300 s duration: (A) Triplicate 1: *H. erinaceus* biofilm;(B) Triplicate 2: *H. erinaceus* biofilm;(C) Triplicate 3: *H. erinaceus* biofilm;(D) Triplicate 4 - nystatin assay (10,000 units/ml);(E) Triplicate 5 - nystatin assay (10,000 units/ml;(F) Triplicate 6 - nystatin assay (10,000 units/ml).

**Figure 6:**
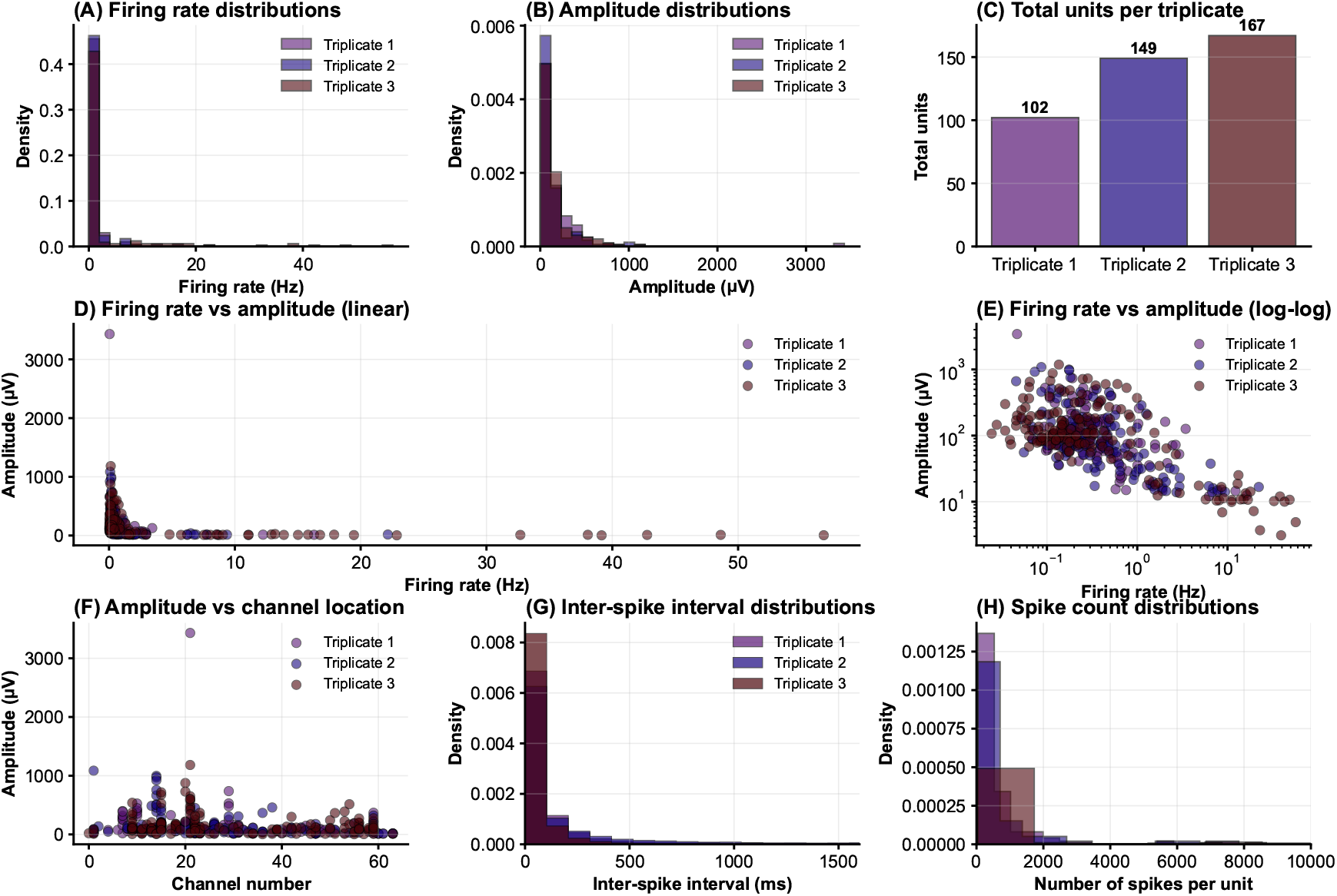
Wild-type discrete unit statistics: (A) Firing rate distribution (Hz);(B) Amplitude distribution (*µ*V); (C) Total units per triplicate (no. units);(D) Firing rate (Hz) vs amplitude (*µ*V); (E) Log-log scale firing rate (Hz) vs amplitude (*µ*V);(F) Amplitude distribution (*µ*V) based on channel location (G) Inter-spike intervals (ms);(H) Spikes per unit.

**Figure 7:**
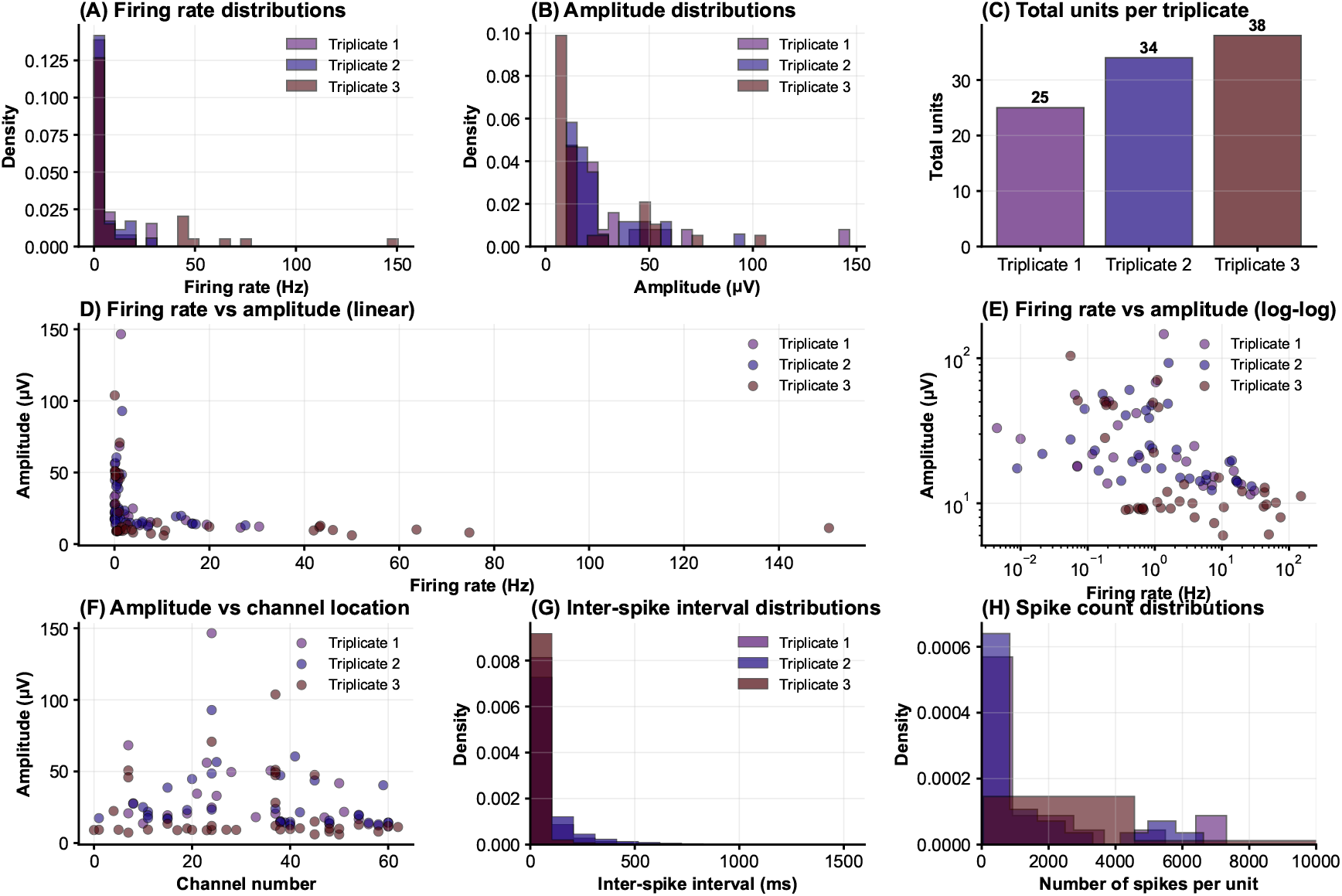
Nystatin assay discrete unit statistics: (A) Firing rate distribution (Hz);(B) Amplitude distribution (*µ*V); (C) Total units per triplicate (no. units);(D) Firing rate (Hz) vs amplitude (*µ*V); (E) Log-log scale firing rate (Hz) vs amplitude (*µ*V);(F) Amplitude distribution (*µ*V) based on channel location (G) Inter-spike intervals (ms);(H) Spikes per unit.

To characterise the effect of nystatin on the extracellular discrete unit spikes we recorded wild-type, nystatin assays and a fungicidal assay. In the wild-type cultures stereotypical spike features were observed across triplicates. Clustered spikes were detected for a total of 418 spiking units (T1: 102 units,T2: 149 units,T3: 167 units). In *H. erinaceus* (*n* = 418), the trough time occurred at 6.04 *±* 0.08 ms, followed by a peak time at 8.73 *±* 0.13 ms, resulting in a trough-to-peak time of 2.68 *±* 0.08 ms. The trough-to-peak time refers, here, to the duration between the maximum negative and maximum positive deflections (post-trough). The waveform width at half-maximum was estimated with a value of 0.81 *±* 0.03 ms. The asymmetry values for the mean waveforms were 0.49 *±* 0.008.

Modification of the transmembrane potential as a result of the application of nystatin was apparent in the lower trough amplitudes of the assayed cultures. In addition the nystatin assay reduced number of units *n* = 97 (T1: 25 units,T2: 34 units,T3: 38 units) total for all triplicates compared with *n* = 418 in the wild-type triplicates. The mean waveforms in the nystatin assay had an estimated trough time of 6.14 *±* 1.59 ms and a peak time of 8.12 *±* 1.89 ms, which produced a trough-to-peak time of 1.97 *±* 0.32 ms. This was 0.71 ms shorter than the average trough-to-peak time of the wild-type cultures. The width at half-maximum was significantly higher at 2.45 *±* 2.45 ms indicating disruption of the stereotypical waveform. The asymmetry values were similar to the wild-type with the nystatin assay exhibiting a mean of 0.53 *±* 0.20. However, higher standard deviations in this assay indicate a more unstable and less stereotypical waveform.

A higher initial positive deflection was also observed in the nystatin cultures resulting in a double humped structured spike morphology for both triplicate 1 and 2. Triplicate 3 showed less deviation from the wild-type cultures with a similar trough-to-peak structure. The spiking units in triplicate 1 and 2 exhibited qualities that would render them borderline them for classification as stereotypical spiking units. The differences in trough-to-peak time between the ionophore nystatin assay and the non-assayed biofilm forming liquid cultures suggest that lipid peroxidation impacted the transmembrane potential. Lipid peroxidation alters the hydrophilicity of the interior of channels in the membrane, which is necessary to transport ions and polar molecules [41]. The modification of the magnitude and morphology of the mean spike waveforms indicates that significant disruption of the plasma membrane occurred and that extracellular electrophysiological spikes, confirmed in the wild-type recordings, were impacted.

To compare the effects of nystatin on the extracellular discrete unit activity with a fungicidal assay we performed triplicate recordings with biofilm forming cultures exposed to 12 % H_2_O_2_ solution. No spiking units or clustered waveforms were detected for the triplicate fungicidal assays. In comparison to the total sorted mean waveform across units and triplicates for the nystatin assay of *n* = 97 and *n* = 418 for the wild types. The resulting waveforms are depicted for a single channel trace and mapped for all channels in Figure 11. Comparison of the frequency components shows no discrete unit activity in the bandpass recordings and electrophysiological activity was confirmed as absent.

Comparing the effects of the fungicidal assay with the nystatin assay suggests that nystatin is a modulator of the transmembrane potential in this species. While cellular leakage occurred in portions of the sample a full mapping was not performed and therefore the effect observed here is likely to be fungistatic rather than fungicidal. The average mean waveform for triplicate 3 of the nystatin assay shows this clearly. While triplicates 1 and 2 provide evidence of fungicidal and fungistatic effects that were localised and varied across the culture. The presence of EPS in all samples is likely to have impeded diffusion of the nystatin solution. As a result it is likely that not all of the physical locations of the spiking units were affected by the nystatin. However, nystatin is also known to target ergosterol and its content in the cell wall varies from species to species with differing levels of effectiveness. For electrophysiological studies in this species, nystatin can be seen as an effective modulator of the transmembrane potential with fungistatic effects rather than a fungicidal assay. A full comparison of the discrete unit waveform statistics is provided in Table 1. The sorted mean waveforms across all units in each triplicate for the wild-type cultures are plotted in Figure 8 (A)-(C). This can be compared with the mean waveforms across all units in each triplicate for the nystatin assay in Figure 8 (A)-(E). Mean waveforms for each of the triplicates in wild-type and nystatin assayed cultures are shown in Figure 9 (A)-(B). The characteristics of the mean waveforms are depicted in Figure 9 (C)-(E). Discrete unit mean waveforms are not provided for the fungicidal assay as the spike sorting was unsuccessful at identifying clusters.

**Table 1:** Comparison of Discrete Unit Waveform Statistics.

| Metric | H. erinaceus (no. units=418) | Nystatin assay (no. units=97) |
| --- | --- | --- |
| Trough time (ms) | $6.04 \pm 0.08$ | $6.14 \pm 1.59$ |
| Peak time (ms) | $8.73 \pm 0.13$ | $8.12 \pm 1.89$ |
| Trough-to-peak time (ms) | $2.68 \pm 0.08$ | $1.97 \pm 0.32$ |
| Width at half-maximum (ms) | $0.81 \pm 0.03$ | $2.71 \pm 2.45$ |
| Waveform asymmetry | $0.49 \pm 0.008$ | $0.53 \pm 0.20$ |

**Figure 8:**
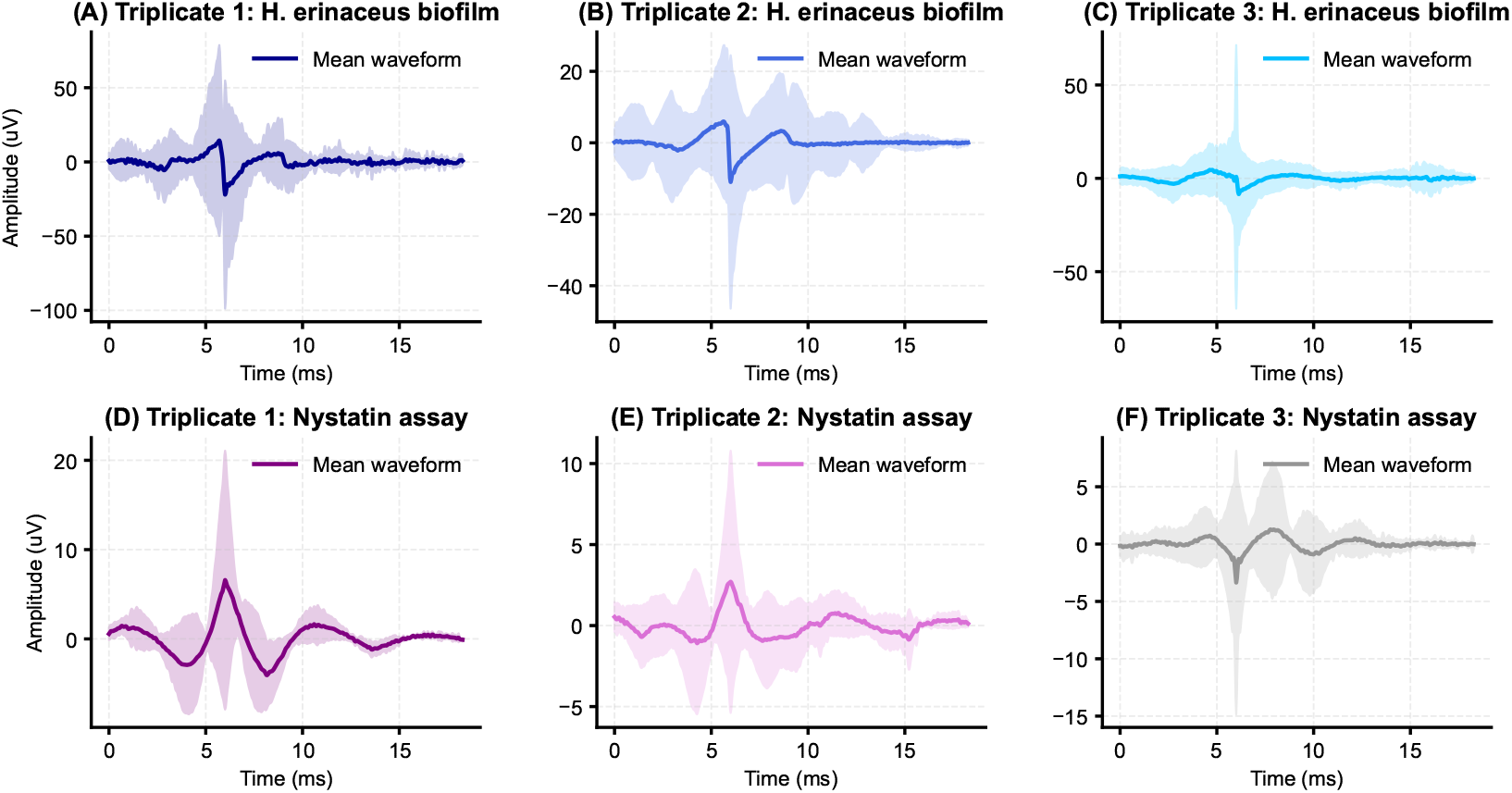
Mean waveforms across all units for: (A) Triplicate 1: *H. erinaceus* biofilm;(B) Triplicate 2: *H. erinaceus* biofilm;(C) Triplicate 3: *H. erinaceus* biofilm;(D) Triplicate 1 - nystatin assay (10,000 units/ml);(E) Triplicate 2 - nystatin assay (10,000 units/ml;(F) Triplicate 3 - nystatin assay (10,000 units/ml.

**Figure 9:**
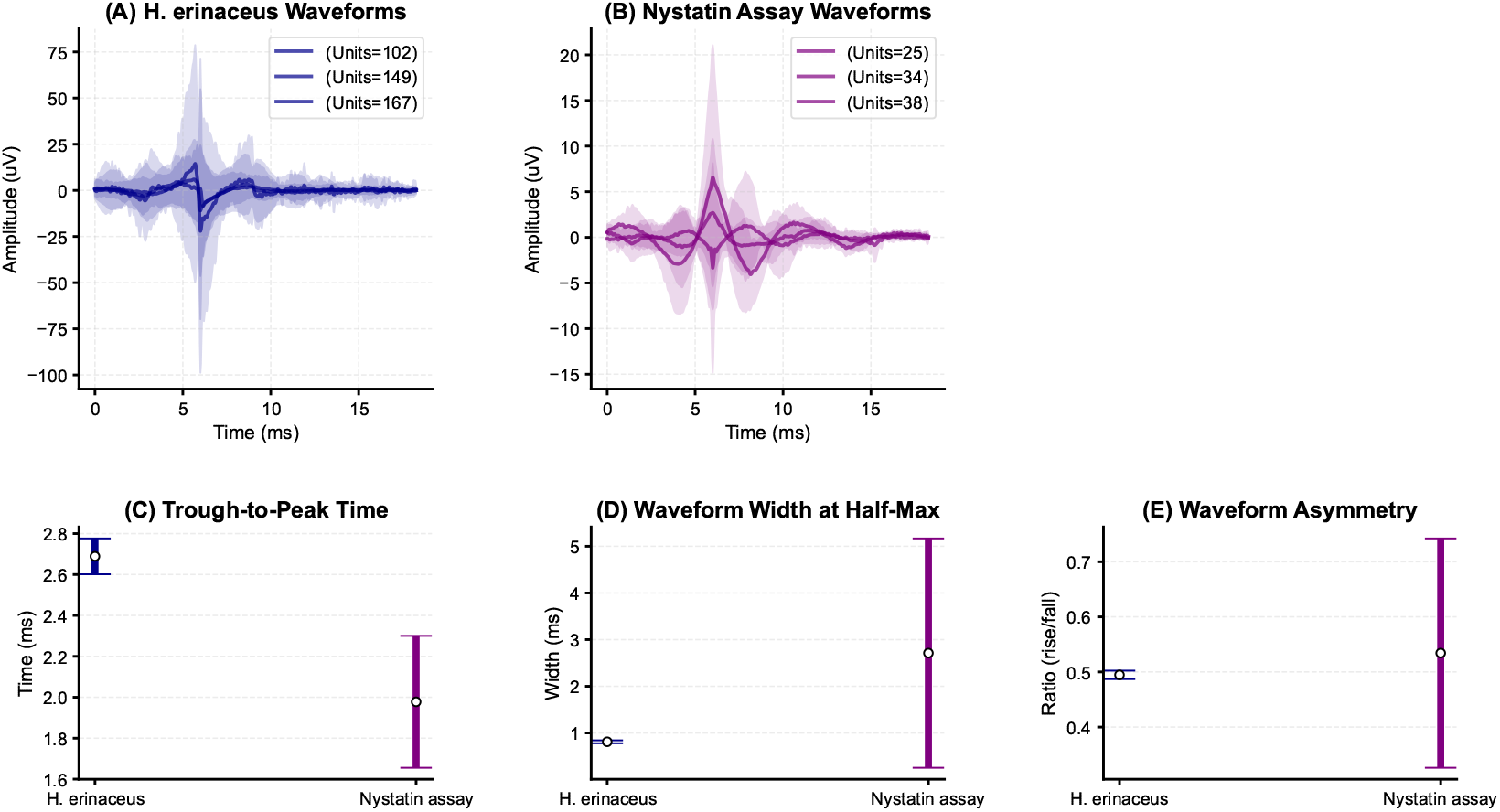
(A) Mean waveforms for triplicates 1-3 *H. erinaceus* biofilm;(B) Mean waveforms for triplicates 1-3: nystatin assay (10,000 units/ml);(C) Comparison of trough-to-peak time for triplicates 1-3 *H. erinaceus* biofilm and nystatin assay (10,000 units/ml);(D) Comparison of width at half-max for triplicates 1-3 *H. erinaceus* biofilm and nystatin assay (10,000 units/ml);(E) Waveform asymmetry triplicates 1-3 *H. erinaceus* biofilm and nystatin assay (10,000 units/ml)

Power spectral density analysis of the full bandwidth of the raw recordings for triplicates combined (see Figure 10 (A)-(C)) showed clear differences between the biofilm, nystatin assay and H_2_O_2_ assay spectra. The dominant frequency in the non-treated biofilm recordings was 7.3 Hz (7.3 Hz (1.08 *×* 10^3^ *µ*V^2^*/*Hz). The dominant frequency in the nystatin assay was 7.3 Hz (2.35 *×*10^2^ *µ*V^2^*/*Hz) and the H_2_O_2_ assay had a dominant frequency of 498.0 Hz (4.96 *×*10^*−*2^ *µ*V^2^*/*Hz). The coefficient of variation in the biofilm spectra was 25.0% compared with 3.4% for the nystatin assay and 15.7% for the H_2_O_2_ assay. Average power was 1.20 *×*10^4^ *µ*V^2^ in the untreated biofilms compared with 2.37 *×*10^3^ *µ*V^2^ in the nystatin assay and 1.38 *×*10^1^ *µ*V^2^ in the H_2_O_2_ assay. The various bandwidths can be seen in more detail in Figure 10 (D)-(G). Of particular note is the presence of narrowband noise in the H_2_O_2_ assay indicating that non-biological sources dominated and the fungicidal assay was successful. The broadband distribution with the exception of minor humps in the viable samples indicates the presence of a broadband signal source typical of electrophysiological rather than EMI or related spikes. The power distribution across frequency bands by triplicate group indicates that the integrated power of both the nystatin and fungicidal assays was lower across all bands compared with the untreated biofilm. The reduction in the power spectra at the lower frequencies (0-500 Hz) in the fungicidal assay was pronounced. The 1*/f* noise exponent in the non-treated biofilm power spectral density was 1.557, with a spectral roll-off of *−*1.6 dB*/*decade. This compares with 1*/f* noise exponent of 0.890 in the nystatin assay and 0.380 in the H_2_O_2_ assay (see Figure 10 (I)). 1*/f* noise exponent analysis therefore indicates that the fungicidal assay was successful for the majority of the area of the sample in the MEA chamber and supports the spike sorting results. Nystatin showed a significantly different 1*/f* noise exponent compared with the non-treated biofilms. However, it was within a feasible range for spontaneous extracellular activity. The spectral roll-off for both assays had a similar pattern: in the nystatin assay *−*0.9 dB*/*decade and *−*0.4 dB*/*decade for the H_2_O_2_ assay. Both power spectral density measures indicate fungicidal action in the H_2_O_2_ assay and spatially limited or fungistatic effects in the nystatin assay. Spatial limitations on diffusion of nystatin solution are likely as a result of the presence of the biofilm. Narrowband spikes in the H_2_O_2_ may be the result of lower magnitude EMI when compared to the baseline electrophysiology.

**Figure 10:**
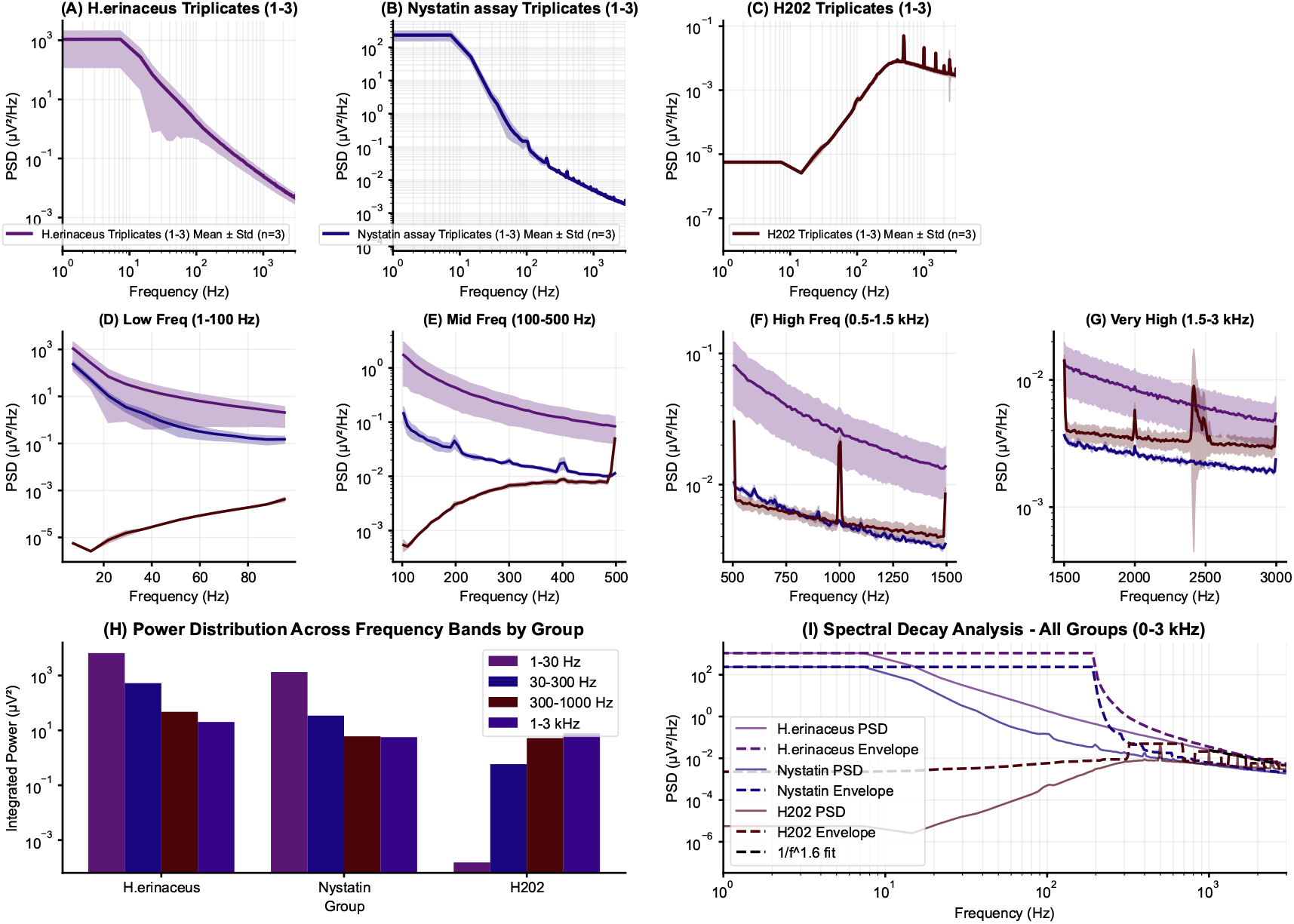
Raw power spectral density for all channels (0.1-3000 Hz): (A) Triplicates 1-3: *H. erinaceus* dispersed;(B)Triplicates 1-3: nystatin assay.;(C)Triplicates 1-3: nystatin assay 12 Hr.;(D) All triplicates comparison of low frequency bands (1-100 Hz);(E)All triplicates comparison of mid frequency bands (100-500 Hz);(F)All triplicates comparison of high frequency bands (0.5-1.5 kHz);(G) All triplicates comparison of highest frequency bands (1.5-3 kHz);(H) Power distribution across frequency bands by triplicate group;(I) Spectral density analysis all groups (0-3 kHz).

**Figure 11:**
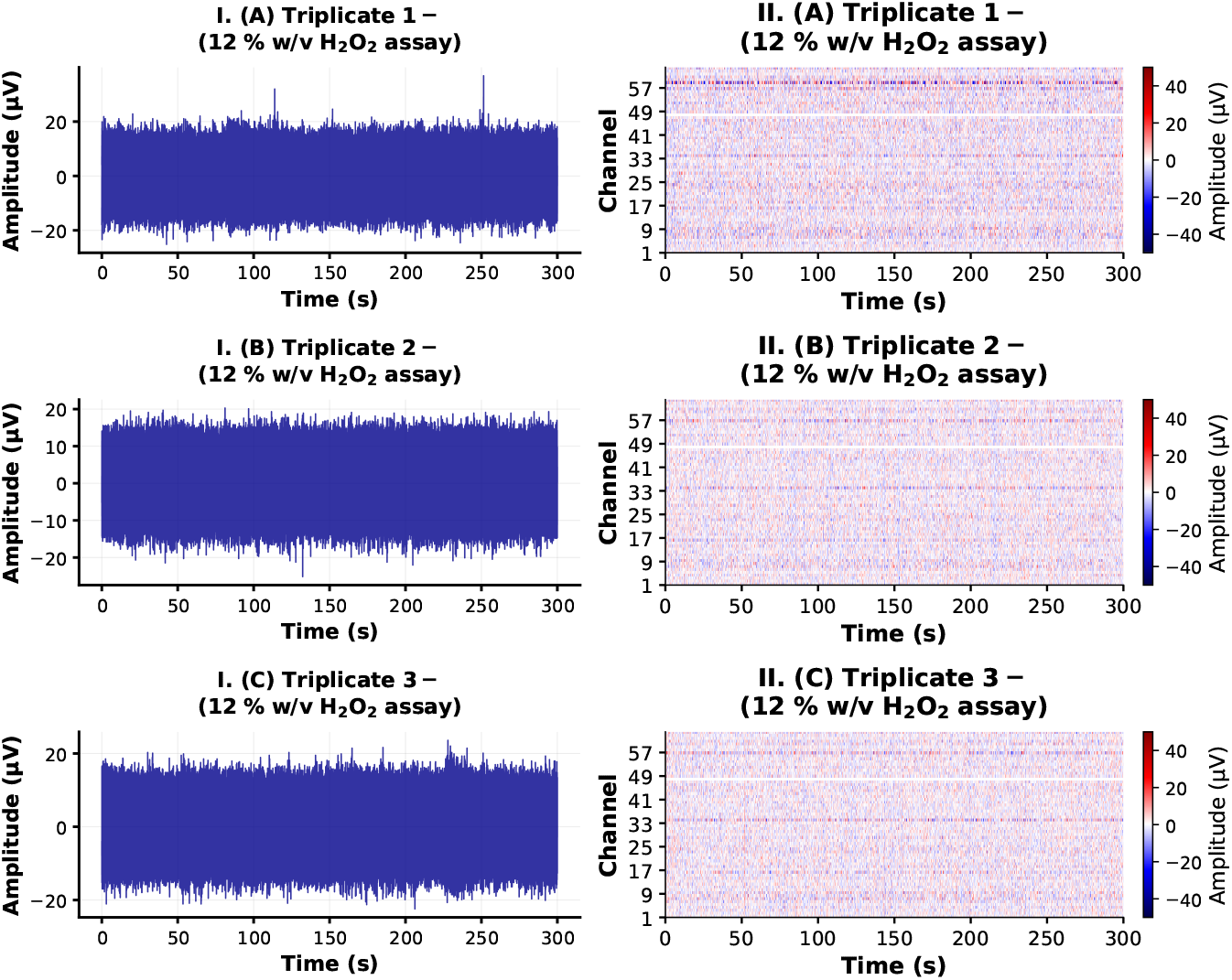
12% w/v H_2_O_2_ solution fungicidal assay triplicates 300 s duration: I. (A) Single channel trace triplicate 1;I. (B) Single channel trace triplicate 2;I. (C) Single channel trace triplicate 3. II. (A) All channels in trace map plot triplicate 1;II. All channels in trace map plot triplicate 2; II. (C) All channels in trace map plot triplicate 3.

## 4. Discussion

In this study, MEAs with hard gold microelectrodes with radius of 100 *µ*m in an array of 64 single ended recording electrodes, including a reference electrode, were used to parse the short-timescale extracellular electrophysiology of liquid biofilm forming cultures of *H. erinaceus* mycelium. The overall system was highly customisable and allowed for arbitrary recording parameters (e.g. spikeinterface) and offline sorting methods (Kilosort4). Following bandpass filtering (150-3000 Hz) discrete unit spikes of milliseconds duration were identified throughout the wild-type recordings. Due to the radius of the microelectrodes, and with reference to environmental scanning micrographs of the in-situ mycelium, the spikes are presumed to originate from sections of approx. *≤* 200 *µ*m of individual hypha that formed clusters on top of the microelectrodes. EPS materials are likely to facilitate electrical and ionic signaling pathways in the wild-type samples and resulted in a large number of total spiking units across triplicates (T1: 102 units,T2: 149 units,T3: 167 units).

Offline spike sorting was implemented using the Kilosort4 sorting algorithm with parameters relevant to the recording setup. Chunks of 900 s were selected from each of the triplicate cultures beginning at 0 s in the recording. All triplicate recordings commenced after 2 hr of resting time in the chamber to allow for recovery from transfer shock. The unusual nature of detected discrete unit spikes means that competing non-biological sources should be examined. Noise, such as 50 Hz, can be discounted due to the grounding and reference procedures in the analog recordings [42]. In addition spike sorting was performed with a high pass filter of 150 Hz which is sufficient to remove lower frequency components. The use of a Faraday cage and RF shielding in addition to mechanical stabilisation limited the impact of external noise.

Donnan potentials are another possible source of noise in recordings of transmembrane potentials using various types of proximal electrodes [43]. In general, Donnan potentials are associated with slow potential shifts that can last for several seconds to minutes and not discrete transients with defined waveforms and millisecond durations [44]. Therefore, these passive potentials can be decisively ruled out in this study. The uniformity of the sorted spike waveforms suggests that active and highly stereotypical spikes were identified and were disrupted by lipid peroxidisation. The putative link between the discrete unit potential and the plasma membrane requires further study. However, the discrete unit activity was modulated significantly by disruption of the cell wall. In contrast, the fungicidal effects on electrophysiology in 12% w/v H_2_O_2_ assay were clear leading to zero spiking units and failed sorting due to no clusters. As a result, electrochemical noise can be distinguished from electrophysiology in these recordings.

The waveform modulating rather than fungicidical effects of nystatin can be explained via biophysical differences in Basidiomycetes cellular composition and EPS components. The multicellular structure, biofilm formation and lower permeability compared to yeasts likely make this species highly tolerant or largely resistant to nystatin. However, ion leakage, stress and disrupted signaling can still occur in the absence of fungicidal effects. As a result, nystatin appears to have a specific role as a transmembrane potential modulator leading to changes in the spike waveform morphology as recorded here via extracellular microelectrodes. There are a number of putative sources for this modulation capacity including the formation of pores in the cell wall and subsequent leakage of small molecules or ions (e.g. K_+_, Ca_2+_, altered membrane fluidity or disrupted membrane protein function (e.g., transporters or signaling proteins). The resulting traces pre-and-post filtering were significantly different comparing the H_2_O_2_ and nystatin assays. These results establish a modulating ionophore and fungicidal electrophysiological cessation assays.

The physiological basis for the discrete extracellular spikes is still not entirely clear. Early research on fungal electrical signaling suggested the role of proton pumps (H^+^) and various ion channels (Ca^2+^, Cl^-^) [6, 45]. The spontaneous potentials in *Neurospora* were attributed to an electrogenic H^+^ pump or altered membrane selectivity. H^+^ and Cl^-^ were determined to be the primary ions that contribute to the inward currents during potential firing [6]. The spontaneous potential-like firing in *A. bulbosa* and *P. ostreatus* exhibited distinct responses to current injection compared to classical animal models [2]. Later patch clamp studies are of more relevance where milliseconds duration spikes in the range of 1-100 ms are visible at approximately 100 times the magnitude of the extracellular spikes observed here post-sorting [13, 14, 46, 7]. It is not entirely surprising that the intracellular potentials recorded from patch clamp and similar methods may have an extracellular correlate given the presence of various relevant ion channels and the transmembrane potential [47]. In theory, the transmembrane current and the extracellular potential follow the same time course, with minor differences due to noise, and are roughly equivalent to the first derivative of the transmembrane potential [2]. Vibrating probe studies are not immediately relevant because of the different recording properties (lock-in amplifiers) and spatial scanning methods [3]. Infra-slow electrical activity is relevant where electrophysiology has been identified as contributing to intermediate and slow waves of electrical activity via pharmacological assays [24]. Glutamic acid was found to increase the frequency of infra-slow oscillations in *H. erinaceus* mycelial biofilms. Nystatin was found to reduce the frequency and completely remove large magnitude macro-scale waves of electrical activity [24]. The development of concurrent infra-slow (*≤* 1 Hz) and intermediate sample rate setups would be useful to determine whether there is a causal relationship between these bioelectrical oscillations and spikes. Continuous signals, e.g. equivalent to local field potentials, at intermediate frequencies (1 - 150 Hz) can also be examined. Different ion and voltage-gated channels have been described in filamentous fungi [47]. Voltage-gated proton channels were related to the fungal Hv channels and conformational changes that resulted in gate opening and proton conduction. Fungal Hvs share 20–29 % sequence identity with the human voltage-gated proton channel hHv1 [48]. Whole-genome sequencing has facilitated the identification of genes likely encoding homologues of K^+^, Ca^2+^, transient receptor potential (Trp), and mitochondrial Ca^2+^ uniporter channels [49]. Homology searches for Basidiomycota have shown presence of putative voltage-gated channels in response to multiple ions (including Ca^2+^, Cl^+^, H^+^ and Na^+^) and the signalling molecule glutamate [50, 51]. The combination of homology searches and ion channel isolating patch clamp studies are useful for identification of these putative voltage-gated channels. However, once the channels have been identified the short duration and non-physiological conditions (i.e. protoplast extrusion) of current patch clamp methods are major limitations in detection of discrete unit potential spikes.

The size of the microelectrodes in the array (radius of 100 *µ*m) is another consideration for the interpretation of results in this study. In principle, if electrodes are too large electrochemical and bulk effects may dominate depending on the characteristics of the overall system. If the electrodes are too small then the signal propagation along the continuous structure of the plasma membrane may be recorded only in part. This complicates the concept of a single spiking unit as adapted from neural recordings [33]. Sections of hyphae may be defined experimentally based on the observation of dolipore septum with each hyphal section defined as a spiking unit or where different electrical polarisation effects are observed in the proximal region. Alternatively, larger continuous structures can be defined relative to the microelectrode size, spacing pitch of the microelectrodes and the overall size of the array. These considerations can be examined algorithmically via multi-parameter spike sorting and clustering. Slightly larger electrodes may have an advantage, in the case where units are continuous structures punctuated every 100-200 *µ*m, due to the nature of signal propagation in the plasma membrane as measured in proximate regions extracellularly. Simultaneous nanoelectrode recordings inside and outside an individual septa or hypha with continuous intracellular structure could prove valuable yet they might damage sections of hyphae or introduce damage specific signaling [52]. In the MEA system outlined in this paper, the multi-unit spiking dynamics are identified for the first time at this timescale and frequency bandpass and allow for comparative investigation of smaller electrode sizes and higher densities. The unique morphological constraints of mycelium should be taken into account and direct inference from existing methods in neural recordings may not transfer. For instance, the physical length scale of spiking units and the interior structure of septate junctions and the plasma membrane.

## 5. Conclusion

In this paper we outlined methods to record and detect discrete unit spikes in biofilm forming *H. erinaceus* liquid cultures using a custom designed hard gold coated microelectrode array. The spike sorting algorithm Kilosort4 was utilised for offline sorting of the discrete unit potentials with a total of 418 individual spiking units detected across all wild-type triplicates with a mean trough-to-peak time of 2.68 *±* 0.08 ms. Nystatin was observed to disrupt the structure of the cell wall and plasma membrane in environmental scanning electron microscopy. There were significant changes in the waveform characteristics (e.g. mean trough-to-peak time of 1.97 *±* 0.32 ms), amplitude of spikes and the number of detected spiking units with a total of 97 across triplicates. The modulating capacity of nystatin in the assayed cultures provided evidence for a putative physiological origin of discrete unit activity in the plasma membrane and associated transmembrane potential. A fungicidal assay utilising 12% w/v H_2_O_2_ solution resulted in zero spiking units confirming physiological and electrophysiological origin of the previously detected spikes. The methods, hardware and signal processing techniques presented in this paper are expected to aid in the development of standardisation of studies of extracellular electrophysiology of mycelium.

## 6. Acknowledgment

The authors would like to thank David Patton for assistance with the environmental scanning electron microscopy. The research has been conducted under the framework of the FUNGATERIA (www.fungateria.eu) project, which has received funding from the European Union’s HORIZON-EIC-2021-PATHFINDER CHALLENGES programme under grant agreement No. 101071145. It is co-funded by the UK Research and Innovation grant No. 10048406.

## 7. Data availability

The datasets used and/or analysed during the current study available from the corresponding author on reasonable request.

